# Using a simple cellular assay to map NES motifs in cancer-related proteins, gain insight into CRM1-mediated NES export, and search for NES-harbouring micropeptides

**DOI:** 10.1101/2020.08.04.234948

**Authors:** Maria Sendino, Miren Josu Omaetxebarria, Gorka Prieto, Jose Antonio Rodriguez

**Author notes:** Correspondence to: Dr. Jose Antonio Rodriguez. Department of Genetics, Physical Anthropology and Animal Physiology, University of the Basque Country (UPV/EHU), Leioa 48940, Spain.

## Abstract

The nuclear export receptor CRM1 (XPO1) recognizes and binds specific sequence motifs termed nuclear export signals (NESs) in cargo proteins. About 200 NES motifs have been identified, but over a thousand human proteins are potential CRM1 cargos, and most of their NESs remain to be identified. On the other hand, the interaction of NES peptides with the “NES-binding groove” of CRM1 has been studied in detail using structural and biochemical analyses, but a better understanding of CRM1 function requires further investigation of how the results from these *in vitro* studies translate into actual NES export in a cellular context. Here we show that a simple cellular assay, based on a recently described reporter (SRV_B/A_), can be applied to identify novel potential NESs motifs, and to obtain relevant information on different aspects of CRM1-mediated NES export. Using cellular assays, we first map 19 new sequence motifs with nuclear export activity in 14 cancer-related proteins that are potential CRM1 cargos. Next, we investigate the effect of mutations in individual NES-binding groove residues, providing further insight into CRM1-mediated NES export. Finally, we extend the search for CRM1-dependent NESs to a recently uncovered, but potentially vast, set of small proteins called micropeptides. By doing so, we report the first NES-harbouring human micropeptides.

## INTRODUCTION

Cellular homeostasis requires a continuous trafficking of proteins between the nucleus and the cytoplasm. For most proteins, nucleocytoplasmic transport is an active process, carried out by a family of soluble receptors termed karyopherins (Pemberton and Paschal, 2005; Tran et al., 2014; Cautain et al., 2015). Transport receptors recognize and bind specific amino acid sequence motifs in the cargo proteins that dictate import into the nucleus (nuclear localization signals or NLSs) or export to the cytoplasm (nuclear export signals or NESs). NLSs are recognized by nuclear import receptors, such as the Importin α/β heterodimer (Soniat and Chook, 2015), whereas NESs are recognized by nuclear export receptors, such as CRM1 (also called XPO1) (Hutten and Kehlenbach, 2007).

According to the number of cargos, CRM1 is the main receptor for protein nuclear export. In addition to several dozens of well-validated cargos, a global proteomics analysis in HeLa cells identified more than 1000 proteins that are probably exported by CRM1, constituting what was termed the “CRM1-dependent nuclear exportome” (Kirli et al., 2015). Binding to CRM1 is usually mediated by short amino acid motifs in the cargo protein, often referred to as “classical” or “leucine-rich” NESs, which show a pattern of 4 or 5 characteristically spaced hydrophobic residues. Hundreds of different NES motifs that differ widely in their amino acid sequence and their affinity for CRM1 (Kutay and Guttinger 2005; Xu et al., 2012; Fu et al., 2013) have been identified. According to the spacing between their hydrophobic residues and the conformation they adopt when bound to CRM1, NESs have been classified into four main groups (class 1, 2, 3 and 4) (Kosugi et al., 2008; Fung et al., 2017). In addition, certain class 1 NESs (so called “minus” motifs) have been reported to bind CRM1 in a reverse orientation, and are thus classified as class 1-R (Fung et al., 2015).

CRM1 is often altered in human tumors. The most common cancer-related CRM1 alteration is protein overexpression, which has been detected in most types of solid and haematological malignancies (reviewed in Sendino et al., 2018). In addition, a recurrent hotspot mutation in CRM1 residue E571 is highly prevalent in specific types of leukaemia and lymphoma (see below). CRM1 is increasingly regarded as an important target for cancer therapy. Therapeutic inhibition of CRM1, mainly using a family of drugs termed selective inhibitors of nuclear export (SINEs) has been extensively validated in preclinical studies, and is being evaluated in multiple clinical trials (Sendino et al., 2018). The most clinically advanced SINE compound, KPT-330 or selinexor (marketed as XPOVIO™) has been recently (July 2019) approved by the FDA for the treatment of relapsed or refractory multiple myeloma in combination with dexamethasone (Syed, 2019). The antitumor activity of CRM1 inhibitors is thought to be mediated, at least in part, by their ability to restore the normal localization and activity of cancer-related CRM1 cargos that are mislocalized in tumors due to altered CRM1 expression or function. To better understand the biological effects of CRM1 inhibitors in cancer patients, it is thus essential to extend our current knowledge of cancer-related CRM1 cargos. In this regard, by combining the “CRM1-dependent nuclear exportome” protein set (Kirli et al., 2015) with the subset of “Cancer-related gene” entries of the Human Protein Atlas (https://www.proteinatlas.org/), we recently proposed the term “XPO1-cancer exportome” (Sendino et al., 2018) to refer to a group of 136 cancer-related proteins that are known or potential CRM1 cargos. Importantly, the NES motifs that could mediate CRM1-dependent export of these proteins remain unknown in most cases. Identifying sequence motifs with nuclear export activity in these proteins would further validate them as CRM1 cargos, and pave the way for subsequent functional analyses.

NES peptides bind to a hydrophobic groove in the surface of CRM1. This NES-binding groove is wider at one end (where the N-terminus of most known NESs binds), and then displays a constriction, becoming narrower at the other end. A series of landmark studies a decade ago showed that NES peptides dock into five hydrophobic pockets of the CRM1 groove, and identified several key amino acids, including I521, L525, F561 and F572 (residue numbering of human CRM1 is used throughout the text) that establish hydrophobic interactions with NES residues (Monecke et al., 2009; Dong et al., 2009a; Güttler et al., 2010). Subsequent analyses have shown that different NES peptides can adopt different backbone conformations when bound to CRM1 (Fung et al., 2017). These studies have also identified K568 as a key residue in CRM1 groove, (Fung et al., 2017). This residue not only contributes to NES binding by hydrogen bonding with the NES backbone, but also appears to function as a “specificity filter” that physically blocks binding of those NES-like peptides whose structural features are not optimal for docking into the groove (Fung et al., 2017). Importantly, K568 establishes electrostatic interactions with E571, the CRM1 residue recurrently mutated in certain haematological malignancies (Puente et al., 2011; reviewed in Sendino et al., 2018). We have previously shown that the cancer-related mutation E571K subtly alters binding of certain NESs (García-Santisteban et al., 2016), and acts as an oncogenic driver (Taylor et al., 2019).

Structural and biochemical data have significantly advanced our understanding of NES binding and export by CRM1, but several questions remain to be explored. For example, combined mutations of multiple hydrophobic residues in CRM1 groove have been shown to disrupt binding (Dong et al., 2009a) and export (García-Santisteban et al., 2016) of a few NESs, but the contribution of individual hydrophobic groove residues to the export of different NES classes has not been investigated. On the other hand, while the effect of K568 and E571 mutations has been separately analysed, a direct comparison of how mutation of these residues affects NES export has not been yet carried out.

In the present study, we apply a recently described nuclear export reporter termed SRV_B/A_ (Figure 1A-C) (Taylor et al., 2019) to map new sequence motifs with nuclear export activity in “XPO1-cancer exportome” proteins, and to gain further insight into CRM1-mediated NES export. In addition, we use this reporter to describe, for the first time, the presence of CRM1-dependent NESs in a recently uncovered and largely uncharacterized set of small proteins termed micropeptides (Hartford and Lal, 2020). Altogether, our work illustrates how simple transfection-based cellular assays can be applied to obtain relevant information on different aspects of CRM1-mediated nuclear export.

**Figure 1.**
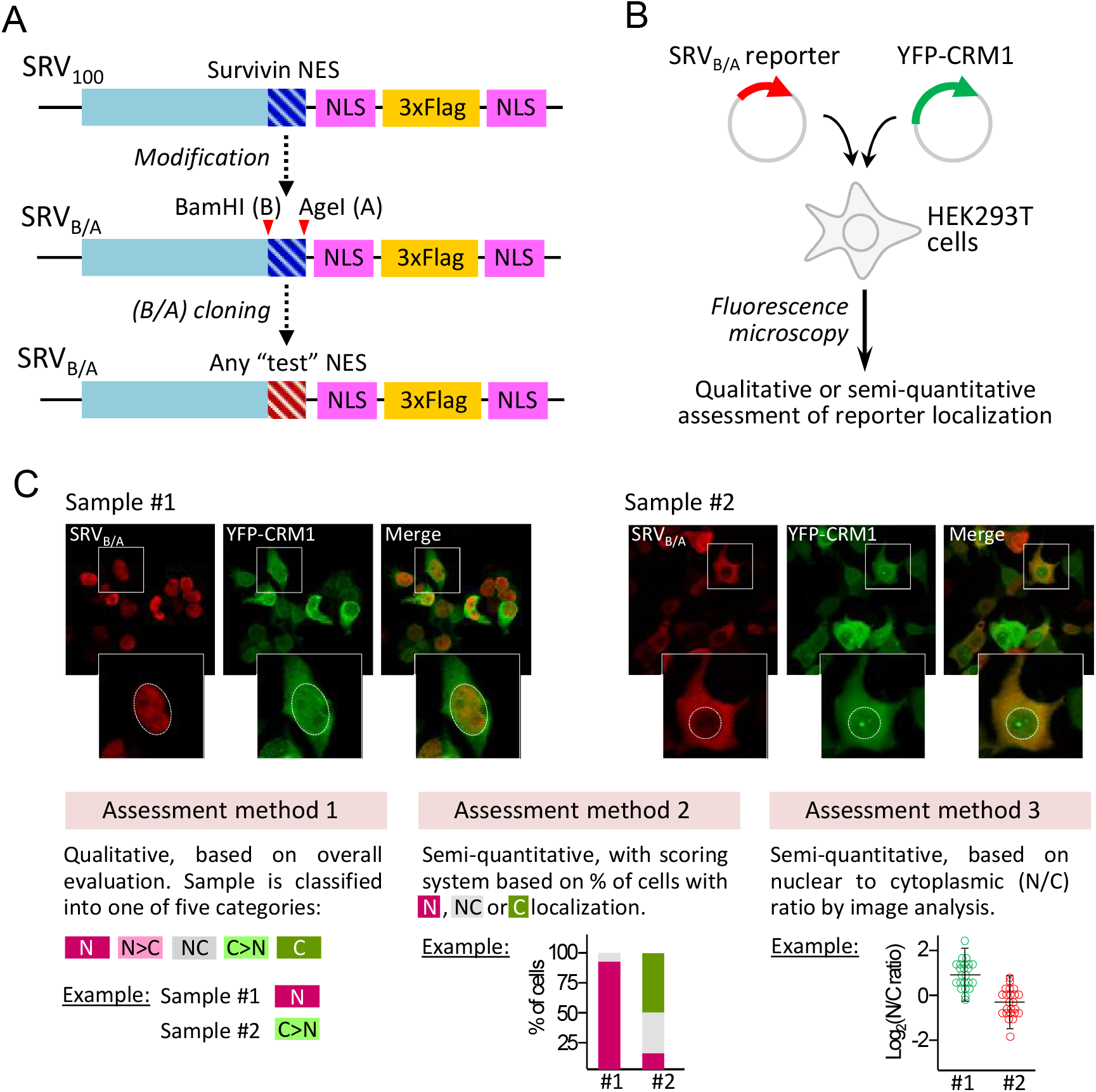
Description and use of the SRV_B/A_ reporter with different assessment methods. *A*. Schematic illustration showing the development and configuration of SRV_B/A_. This reporter is derived from the previously described SRV100 reporter containing the NES of survivin (García-Santisteban et al., 2016). As originally described in a recent report (Taylor et al., 2019), the plasmid encoding SRV100 was modified to introduce two novel restriction sites (BamHI/AgeI), which allows replacing survivin NES by any other NES motif. Besides a survivin amino-terminal fragment and the cloned NES motif, the SRV_B/A_ reporter includes two copies of the SV40 NLS (PKKKRKV) separated by three tandem copies of the Flag epitope. *B*. To carry out SRV_B/A_-based assays, HEK293T cells are co-transfected with plasmids encoding the reporter and the desired YFP-CRM1 variant. After anti-Flag immunostaining the localization of the reporter is determined using fluorescence microscopy. *C*. Microscopy images show examples of the results of the assay with two reporters that contain different NES motifs. Insets show the magnified image of a single cell of each sample, with the nucleus delimited by a dotted line. Under the images, the three different methods used to manually assess the localization of the reporters in this study are described. Method 1 is a qualitative assessment, where the sample is ascribed, according to overall reporter localization, to one of five categories: exclusively nuclear (N), mainly nuclear (N>C), nuclear and cytoplasmic (NC), mainly cytoplasmic (C>N) or exclusively cytoplasmic (C). In the example, sample#1 is classified as N, and sample#2 as C>N. Method 2 is a more detailed, semiquantitative assessment that we have previously used with the SRV100 reporter (García-Santisteban et al., 2016). The localization of the reporter in at least 200 individual cells per sample is evaluated, and classified as exclusively/mainly nuclear (N), nuclear and cytoplasmic (NC), or exclusively/mainly cytoplasmic (C). Based on the percentage of cells showing N, NC or C localization of the reporter, a nuclear export score (termed “SRV export score”) between 0 (no export) and 100 (complete export) is calculated as described in Methods. Graphs show the results for sample#1 and sample#2, corresponding to “SRV export scores” of 4 and 70, respectively. Method 3, the most detailed and laborious assessment method used here, is based on image analysis to determine the nuclear to cytoplasmic (N/C) ratio of the Alexa Fluor 594 fluorescent signal corresponding to the SRV_B/A_ reporter in an average of 50 cells per sample. Graphs illustrating the results for sample#1 and sample#2 are shown.

## MATERIALS AND METHODS

### *In silico* NES prediction

From the putative “XPO1-cancer exportome” (Sendino et al., 2018), we selected 112 proteins that had been classified as CRM1 cargo A or cargo B by Kirli et al. (2015). The amino acid sequence of these proteins was analysed using two different bioinformatics tools: Wregex (Prieto et al., 2014) with the recommended configuration and NESmapper (Kosugi et al., 2014) with a minscore of 0 and trained profile. It has been reported that some NES motifs (so called “minus” motifs) can bind CRM1 in an opposite orientation to that of most previously described motifs (hereafter referred to as “plus” motifs) (Fung et al., 2015). In order to predict potential “minus” motifs, the amino acid sequence of each protein was reversed before being entered as input for the analysis. *In silico* predicted NES motifs were ranked according to the score provided by Wregex and NESmapper. Rank 1 motifs were defined as those included in the first quartile of both programs. Rank 2 motifs were defined as those included in the first quartile of one of the programs and the second quartile of the other. This ranking was the criterion used to select those predicted candidate NES (cNES) motifs to be experimentally tested. Micropeptide sequences, on the other hand, were obtained from the high confidence set of the SmProt small protein database (Hao et al., 2018) and analysed using Wregex and NESmapper. A set of 10 candidates in the first quartile of both programs was selected for experimental testing. The analysis was limited to the prediction of “plus” motifs.

### Plasmids, cloning procedures, and site-directed mutagenesis

The plasmid encoding the Rev(1.4)-GFP reporter (Henderson and Eleftheriou, 2000) was a gift from Dr. Beric Henderson (University of Sydney, Australia). The plasmid encoding the SRV_B/A_ reporter has been recently described (Taylor et al., 2019). This plasmid, derived from SRV100 (García-Santisteban et al., 2016), allows cloning of desired cNES motifs as BamHI/AgeI fragments. These cloning sites and the reading frame in SRV_B/A_ allow easily shuttling cNES-coding DNA sequences to and from the Rev(1.4)-GFP plasmid. DNA sequences encoding cNES motifs were purchased as gBlocks (IDT), digested with BamHI and AgeI and cloned into Rev(1.4)-GFP and SRV_B/A_ plasmids.

The plasmids encoding YFP-CRM1 wild type, and several mutants used in this study (4X, F572A, E571K and A541K) have been previously described (Rodríguez and Henderson, 2000; García-Santisteban et al., 2016). The remaining CRM1 mutants (I521A, L525A, F561A and K568A) were generated by site directed mutagenesis using the Quick-Change Lightning Site-Directed Mutagenesis Kit (Agilent Technologies) according to the manufacturer’s instructions.

All the new constructs were subjected to DNA sequencing (StabVida) and the absence of any unwanted mutation was confirmed.

### Cell culture, plasmid transfection and LMB treatment

HeLa and human embryonic kidney 293T (HEK293T) cells were grown in Dulbecco’s modified Eagle’s medium (DMEM) supplemented with 10% fetal bovine serum (FBS), 100 U/ml penicillin and 100 μg/ml streptomycin (all from Life Technologies) at 37°C in a humidified atmosphere containing 5% CO_2_. 24 hours before transfection, cells were seeded in 12-well plates with glass coverslips. Plasmid transfections were carried out using X-tremeGENE 9 DNA transfection reagent (Roche Diagnostics) following manufacturer’s instructions. The CRM1 inhibitor Leptomycin B (Apollo Scientific) was used at a final concentration of 30 ng/ml for 3 h.

### Rev(1.4)-GFP and SRV_B/A_ nuclear export assays

The Rev(1.4)-GFP nuclear export assay was carried out as previously described (Henderson and Eleftheriou, 2000). Briefly, Rev(1.4)-GFP plasmids with the different cNES motifs were transfected into HeLa cells. The empty Rev(1.4)-GFP plasmid was used as negative control. Each plasmid was transfected in two wells. 24 hours after transfection, 10 μg/ml cycloheximide and 5 μg/ml actinomycin D (both drugs from Sigma-Aldrich) were added to one of the wells. Only cycloheximide was added to the second well. As detailed in the original report (Henderson and Eleftheriou, 2000), cycloheximide is added to ensure that the fluorescent signal in the cytoplasm corresponds to exported and not to newly synthesized GFP-tagged proteins, while actinomycin D facilitates detection of weaker NESs. After 3 hours of treatment, cells were fixed with 3.7% formaldehyde (Sigma-Aldrich) in phosphate-buffered saline (PBS) for 30 min, washed with PBS, and directly mounted onto microscope slides using Vectashield mounting medium containing DAPI (Vector Laboratories). Samples were analysed using a Zeiss Axioscope fluorescence microscope. To ensure unbiased assessment, the identity of the samples was masked before the analysis. At least 200 transfected cells per sample were examined to establish the proportion of cells where the reporter shows nuclear (N), nuclear and cytoplasmic (NC) or cytoplasmic (C) localization. Based on this proportion, each of the tested motifs was assigned a nuclear export activity score between 0 (no export activity, inactive motif) and 9 (maximal export activity) using the assay scoring system (Henderson and Eleftheriou, 2000).

The SRV_B/A_ assay was carried out in HEK293T cells. Plasmids encoding SRV_B/A_ reporters with the different cNES motifs were transfected alone or co-transfected with the indicated YFP-CRM1 plasmids. 24 h after transfection, cells were fixed with 3.7% formaldehyde (Sigma-Aldrich) in phosphate-buffered saline (PBS) for 30 min, permeabilised with 0.2% Triton X-100 (Sigma-Aldrich) in PBS for 10 min, and blocked in 3% bovine serum albumin (BSA; Millipore) in PBS for 1 h. Cells were incubated with anti-Flag M2 mouse monoclonal antibody (Sigma-Aldrich) diluted 1:400 in blocking solution for 1 h to detect SRV_B/A_ reporters. After washing with PBS, cells were incubated with Alexa Fluor 594-conjugated anti-mouse secondary antibody (Invitrogen) diluted 1:400 for 1 h. Finally, samples were washed and mounted onto microscope slides using Vectashield mounting medium containing DAPI (Vector Laboratories).

Samples were examined using either Zeiss Axioskop or Zeiss Apotome2 fluorescence microscopes, and the results of the assays were assessed using three different methods, illustrated in Figure 1C. Irrespective of the assessment method used, samples were coded before the analysis to avoid potential bias and, although we did not carry out a detailed quantification of YFP fluorescence, we confirmed that the level of expression of YFP-CRM1 plasmids was comparable among the different samples.

In method 1, samples were observed using a Zeiss Axioskop microscope and the overall localization of the reporter was classified as exclusively nuclear (N), mainly nuclear (N>C), equally nuclear and cytoplasmic (NC), mainly cytoplasmic (C>N) or exclusively cytoplasmic (C). A subset of the samples was independently analysed by two of the authors (M.S and J.A.R), and the classification by the two observers was found to be highly concordant. In method 2, previously used with SRV100 (García-Santisteban et al., 2016), the localization of the reporter was determined in at least 200 individual transfected cells per sample using a Zeiss Axioskop microscope. Based on the percentage of cells with nuclear (N), nuclear and cytoplasmic (NC) or cytoplasmic (C) localization of the reporter, a nuclear export score (termed “SRV export score” and ranging between 0 and 100) was derived using the formula 0(%N)+0.5(%NC)+1(%C). The score values for the different reporter/CRM1 combinations tested were represented as a heatmap using the resources available at the Heatmapper web server (www.heatmapper.ca). In method 3, image analysis was used to quantify the intensity of the Alexa Fluor 594 fluorescent signal (corresponding to SRV_B/A_ reporters) in the nucleus and the cytoplasm on an average of 50 individual cells per sample. To this end, optical sectioning images (channels with SRV_B/A_ reporters and YFP-CRM1) or conventional fluorescence images (DAPI-stained nuclei) of 10-15 different areas in each sample were taken using a Zeiss Apotome2 microscope and Zen2.6 Blue edition software. Composite images were created using Fiji software (Schindelin et al., 2012), and analyzed using an ad-hoc script developed previously (Olazabal-Herrero et al., 2019) to automatically quantify the fluorescence intensity in nuclear and cytoplasmic regions. Finally, the nuclear to cytoplasmic (N/C) ratios were calculated, plotted in logarithmic base 2, and samples were compared using the Mann-Whitney U test (GraphPad Prism 7 software). Differences were considered statistically significant when p < 0.05.

### Correlation analyses

In order to statistically test their correlation with the results of Rev(1.4)-GFP assays, the qualitative localization data (N, N>C, NC, C>N or C) obtained using method 1 to assess SRV_B/A_ assays were assigned numerical values. These values (N=0, N>C=1, NC=2, C>N=3 and C=4) were plotted against the Rev(1.4)-GFP assay scores (values between 0 and 9), and the correlation between both sets of data was calculated using the Pearson correlation coefficient. The statistical analysis was performed using GraphPad Prism7 software.

For a subset of well-characterized NES motifs, we tested the correlation between their nuclear export activity in SRV_B/A_ assays and their affinity for CRM1 *in vitro*. To this end, the SRV score value of each NES motif expressed alone (obtained using assessment method 2) was plotted against the previously reported K_d_ value of each CRM1/NES interaction (Fu et al., 2018) and the correlation between both sets of data was calculated as above.

## RESULTS

### Using the SRV_B/A_ assay to identify novel NES motifs in cancer-related CRM1 cargos

Cellular assays based on the localization of reporter proteins are a common approach to evaluate nuclear export activity of putative NES motifs. A prominent example is the Rev(1.4)-GFP nuclear export assay (Henderson and Eleftheriou, 2000). In this widely used assay (cited 330 times, according to the Scopus citation database accessed in May 2020), export activity of candidate NESs is determined by their ability to induce cytoplasmic relocation of the otherwise nuclear Rev(1.4)-GFP reporter. We have previously shown that another cellular assay, based on a reporter termed SRV100, can be used to compare the export activity of different CRM1 variants (García-Santisteban et al., 2016). The original SRV100 reporter contained the NES of survivin but, as recently described (Taylor et al., 2019), we have subsequently modified the plasmid encoding SRV100 to introduce two novel restriction sites (BamHI/AgeI) that allow replacing survivin NES by any other NES (Figure 1A). Importantly, this modified reporter, called SRV_B/A_, was designed in such a manner that candidate NES motifs can be easily shuttled to and from the Rev(1.4)-GFP reporter (Figure 2A).

**Figure 2.**
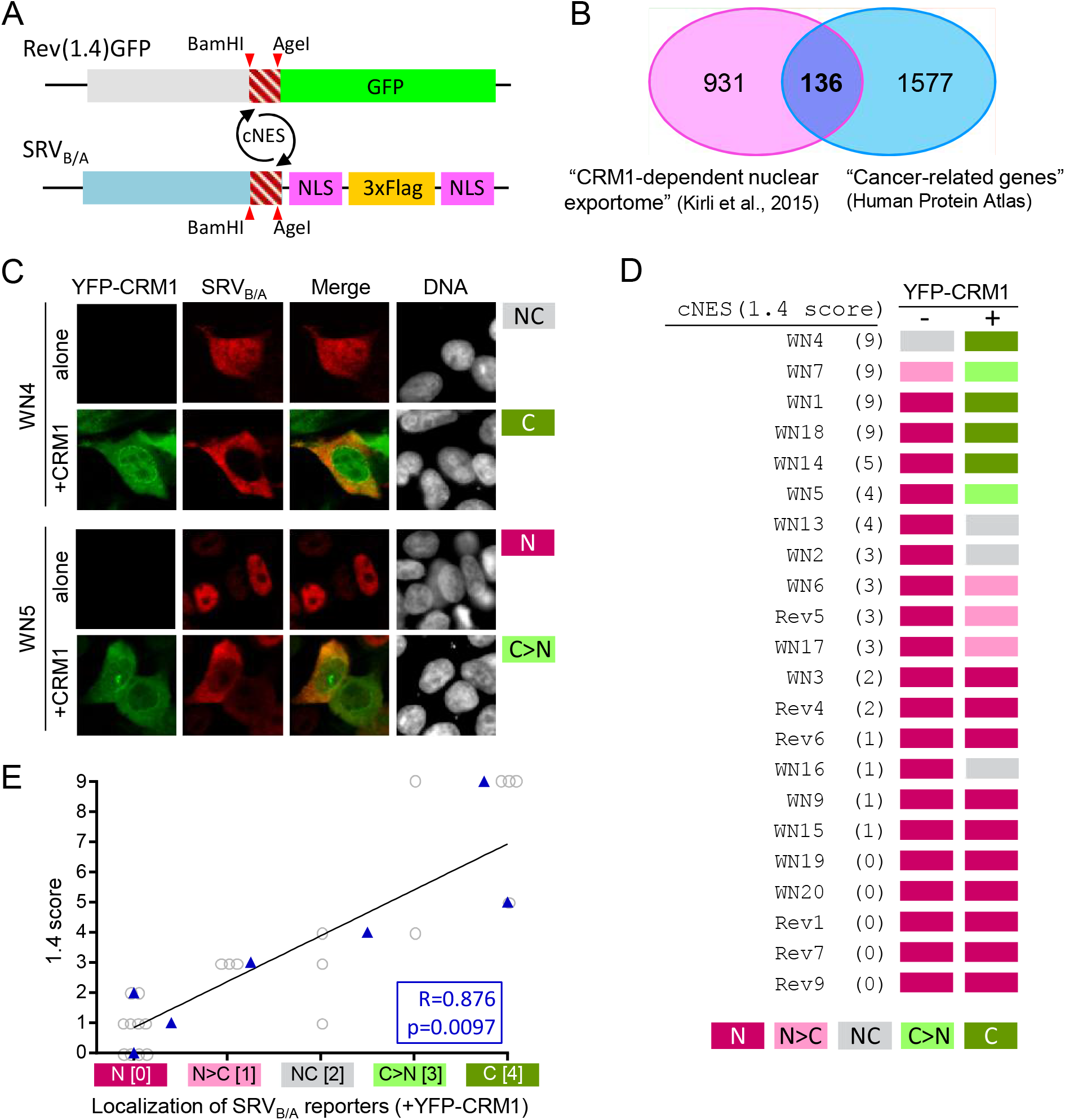
Using the SRV_B/A_ assay to identify novel NES motifs in cancer-related CRM1 cargos. *A*. Schematic representation of Rev(1.4)-GFP and SRV_B/A_ reporters, illustrating how candidate NES motifs (cNES) can be easily shuttled between both reporters by BamHI/AgeI subcloning. *B*. Venn diagram (modified from Sendino et al., 2018) showing the overlap between the list of potential CRM1 cargos reported by Kirli et al (2015) and the group of “Cancer-related gene” defined in the Human Protein Atlas. The 136 common proteins represent what we refer to as the “XPO1-cancer exportome”. *C*. Fluorescence microscopy images showing representative examples of the localization of SRV_B/A_ reporters containing two different cNES motifs (WN4 and WN5, see Supplementary Table 1), when transfected alone or co-transfected with YFP-CRM1 wild type (+CRM1) into HEK293T cells. The DNA-staining dye DAPI was used to visualize the nuclei. As indicated to the right of the images, using assessment method 1 the localization of SRV-WN4 was classified as NC (alone) and C (+CRM1), while the localization of SRV-WN5 reporter was classified as N (alone) and C>N (+CRM1). *D*. Summary of the localization of 22 SRV_B/A_ reporters containing cNES motifs predicted in cancer-related proteins. The cNES ID (Supplementary Table 1) and the 1.4 score for each motif are indicated to the left. The localization of each reporter expressed alone (-) or co-expressed with YFP-CRM1 wild type (+) was assessed using method 1. Ten motifs showed exclusively nuclear (N) localization when co-expressed with YFP-CRM1, and were thus considered inactive in this assay. *E*. Graph showing the correlation between the results obtained with the Rev(1.4)-GFP and the SRV_B/A_ assays for 22 cNES motifs that were tested using both systems. For each motif, the score assigned using the Rev(1.4)-GFP assay (1.4 score, between 0 and 9) was plotted against the localization (N, N>C, NC, C>N, or C) of the corresponding SRV_B/A_ reporter when co-expressed with YFP-CRM1. In order to calculate the correlation coefficient between both sets of data, qualitative descriptions of SRV_B/A_ localizations were assigned a numerical value (N= 0; N>C= 1; NC= 2; C>N= 3; C= 4). The mean of these values for cNES motifs with a given 1.4 score is represented as a blue triangle. Pearson correlation coefficient and p value are indicated.

The SRV_B/A_ assay has not yet been applied to identify novel NES motifs. In order to determine how this assay compares to the well-established Rev(1.4)-GFP assay in this task, we decided to use both systems in an effort to map novel NESs in the proteins that conform the “XPO1-cancer exportome” (Figure 2B). As a first step, we carried out an *in-silico* prediction of putative NES motifs in 112 of these proteins. Their amino acid sequence was analysed with two different programs: Wregex (Prieto et al., 2014) and NESmapper (Kosugi et al., 2014). In order to predict potential reverse (“minus”) NES motifs, the amino acid sequence of each protein was inverted prior to being used as input for the analysis. A ranking approach, based on the score assigned by the programs to each predicted NES, was applied to select a reasonable number of candidate motifs to be experimentally tested. Thus, predicted NES motifs with scores within the first quartile for both programs were designed as Rank 1 candidates. Predicted motifs within the first quartile for one of the programs and within the second quartile for the other were designed as Rank 2. This *in-silico* analysis identified 7 Rank 1 “plus” and 10 Rank 1 “minus” candidates.

The Rev(1.4)-GFP assay was used first to evaluate export activity of all Rank 1 motifs. In addition, all predicted Rank 2 “plus” candidates (19 motifs) were also included in the analysis. All in all, 35 candidates were tested and assigned an export score (hereafter referred to as “1.4 score”) in a range between 0 and 9 (Henderson and Eleftheriou, 2000). As summarized in Supplementary Table 1, 25 of the 36 candidate sequences tested positive in the Rev(1.4)-GFP assay. Representative examples are shown in Supplementary Figure 1A. 19 of these sequences represent novel NES motifs (Supplementary Figure 1B), while six had been previously reported (Fukuda et al., 1996; Toyoshima et al., 1998; Vielhaber et al., 2001; Macchi et al., 2003; Bachmann et al., 2006; North and Verdin, 2007). Among “plus” candidates, 19 out of 25 motifs tested positive, although 3 of them showed borderline activity (1.4 score= 1). On the other hand, 6 out of 10 “minus” candidates tested positive, 4 of them with borderline activity. The mean 1.4 score was 3.84 for “plus” motifs and 1.5 for “minus” motifs (Supplementary Figure 1C).

Next, a subset of 22 motifs with different 1.4 scores were subcloned into the SRV_B/A_ plasmid. These reporters were transfected into HEK293T cells either alone or with YFP-CRM1 and, after anti-Flag immunostaining, their localization was globally assessed by fluorescence microscopy and classified as exclusively nuclear (N), mainly nuclear (N>C), nuclear and cytoplasmic (NC), mainly cytoplasmic (C>N) or exclusively cytoplasmic (C) (assessment method 1, detailed in Figure 1). Representative examples of the localization of two reporters are shown in Figure 2C, and the results obtained with the 22 reporters are summarized in Figure 2D. When transfected alone, all but two reporters (those containing cNES motifs WN4 and WN7, both with a 1.4 score of 9) showed exclusively nuclear localization. On the other hand, when co-transfected with YFP-CRM1, all reporters containing cNES motifs with 1.4 score above 2 showed partial or complete relocation to the cytoplasm. In contrast, all reporters containing cNES motifs with a 1.4 score equal or lower than 2, except SRV-WN16, were classified as exclusively nuclear, even when co-expressed with CRM1. In an attempt to evaluate the correlation between the results obtained with the two assays, 1.4 scores were plotted against the localization of the SRV_B/A_ reporters (when co-transfected with YFP-CRM1). In order to calculate the correlation coefficient, the different SRV_B/A_ localizations were assigned numerical values (N= 0; N>C= 1; NC= 2; C>N= 3; C= 4). As shown in Figure 2E, the results obtained with the Rev(1.4)-GFP and the SRV_B/A_ assays were significantly correlated (R=0.876; p=0.0097). We noted that some sequence motifs with the lowest nuclear export activity (1.4 score equal or lower than 2) may be missed in the SRV_B/A_ assay, when the localization of reporters is qualitatively analysed. Conversely, the SRV_B/A_ assay allowed to detect differences in activity between strong NES motifs. Thus, WN1, WN4, WN7 and WN18 were all assigned a 1.4 score of 9 (the highest possible in this assay), but only SRV-WN4 and SRV-WN7 reporters showed partial cytoplasmic localization when transfected alone, suggesting that WN4 and WN7 motifs are stronger NESs than WN1 and WN18.

In summary, we have experimentally validated 25 sequence motifs with nuclear export activity in “XPO1-cancer exportome” proteins, 19 of which represent novel “plus” or “minus” potential NESs not previously described. Furthermore, these results validate the use of the SRV_B/A_ reporter as a tool to search for novel NESs.

### Using the SRV_B/A_ assay to gain further insight into CRM1-mediated NES export. (i) Effect of single-residue NES-binding groove mutations

The structural analyses that revealed how NES peptides dock into the NES-binding groove of CRM1 were supported by *in vitro* pull-down assays to evaluate how groove mutations affected NES interaction. Thus, it was shown that replacing groove residue A541 with a bulkier lysine (A451K) severely disrupted CRM1/NES interaction (Güttler et al., 2010). A similar disruption was observed when four groove residues that establish hydrophobic interactions with the NES (I521, L525, F561 and F572) were simultaneously mutated to alanine (I521A/L525A/F561A/F572A, hereafter referred to as 4X) (Dong et al., 2009a). Subsequent structural and biochemical studies have identified a key residue in the NES-binding groove, K568, which contributes to NES binding, and may also block docking of non-functional NES-like peptides (Fung et al., 2017). Using the SRV100 reporter-based cellular assay, we have previously confirmed that the A541K and 4X mutations prevent nuclear export of survivin NES (García-Santisteban et al., 2016).

Here we sought to investigate, using the SRV_B/A_ reporter, to what extent individual groove residues contribute to the export of different NES motifs in a cellular context. To this end, we generated a panel of YFP-CRM1 variants bearing single-residue mutations in five groove amino acids (I521A, L525A, F561A, F572A and K568A) whose position is illustrated in Figure 3A. The export activity of these mutants was interrogated using a panel of 14 extensively characterized NES motifs (PKI, superPKI, PAX, HDAC5, FMRP, FMRP-1b, SNUPN, Rev, SMAD4, mDia2, CDC7, X11L2, CPEB4 and hRio2). The binding of these NES peptides to CRM1 has been previously studied using structural and biochemical analyses (Fung et al., 2017), and their export activity has been analysed using a cellular assay (Fu et al., 2018). These well-studied NESs provide a unique resource to evaluate the export activity of the different CRM1 mutants against a variety of motifs that belong to different NES classes (1, 2, 3, 4 and 1-R), and dock into CRM1 groove using different backbone conformations (Fung et al., 2017).

**Figure 3.**
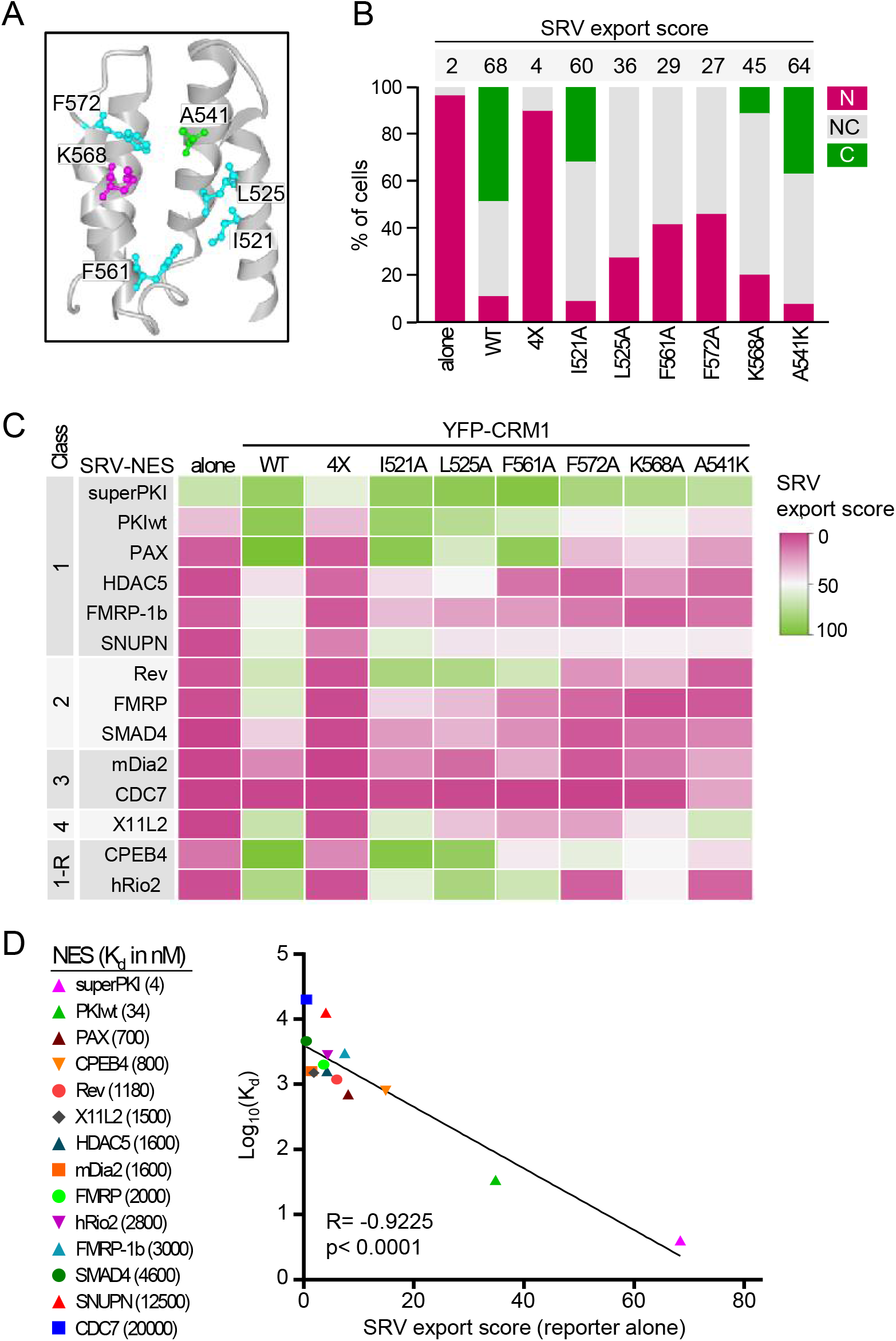
Using the SRV_B/A_ assay to test the effect of single-residue mutations in CRM1 NES-binding groove. *A*. View of CRM1 NES-binding groove generated with NCBI iCn3D viewer from PDB structure 3GJX (Monecke et al., 2009). The residues individually mutated in this study are highlighted using ball and stick representation, while the remaining residues are represented using ribbon style. *B*. Graph shows an example of the results of SRV_B/A_ assays testing export activity of the different YFP-CRM1 mutants against a reporter containing one of the previously-characterized NES motifs (in this case SRV-X11L2). The localization of the reporter was determined in at least 200 cells per sample. Bar colours represent the percentage of cells showing the indicated reporter localization (N, NC or C). The number above each bar indicates the corresponding “SRV export score”, derived as described in Methods section. *C*. Heat map summarizing the results obtained with the 14 reporters (graphs for each reporter are provided in Supplementary Figure 2). The colour indicates the SRV export score for each NES/variant combination, ranging from 0 (no export) to 100 (full export). D. Graph showing the correlation between the previously reported CRM1 binding affinity (expressed as Log_10_(K_d_)) of each NES peptide (Fu et al., 2018) and the SRV export score of the corresponding reporter, when expressed alone. Pearson correlation coefficient and p value are indicated.

SRV_B/A_ reporters containing each of these 14 NES motifs were either transfected alone or co-transfected with YFP-CRM1-encoding plasmids into HEK293T cells. Besides the five mutants indicated above, wild-type YFP-CRM1 and the previously studied A541K and 4X mutants were included in the analysis. All in all, 112 different combinations of SRV-NES reporter/CRM1 mutant (plus each reporter alone) were tested in these experiments. The percentage of cells were the reporter was located exclusively/mainly in the nucleus (N), the cytoplasm (C), or was similarly distributed between nucleus and cytoplasm (NC) was determined by counting at least 200 cells per sample (assessment method 2, detailed in Figure 1). As a representative example, the results obtained with the SRV_B/A_ reporter containing X11L2 NES are shown in Figure 3B, and the graphs for all the NES reporters are presented in Supplementary Figure 2. From these semi-quantitative data, a nuclear export score (hereafter referred to “SRV export score”), ranging between 0 (no export) and 100 (complete export) was derived, as detailed in the Methods section. The SRV scores for the full set of experiments are represented as a heat map in Figure 3C.

When expressed alone, the SRV export score was lower than 10 (corresponding to mainly nuclear localization) for all the reporters except those containing PKI, superPKI and CPEB4 NES motifs. Co-expression with wild-type YFP-CRM1 readily induced nuclear export of all the reporters, with the exception of SRV-CDC7. Simultaneous mutations in the four hydrophobic residues (4X) fully abrogated YFP-CRM1-induced export of all NESs, while individual mutations of these amino acids had less dramatic consequences, as expected. Interestingly, the different single-residue mutations decreased export to a different extent, with I521A and L525A having the mildest effect, and F572A being consistently the most detrimental. These findings suggest that the degree of contribution of these residues to NES export could be expressed as I521=L525<F561A<F572A, irrespective of NES class. Of note, the effect of the K568A mutation was remarkably similar to the effect of F572A, even if these CRM1 residues engage in different types of chemical interaction with the NES. Finally, the A541K mutation severely reduced export of most reporters, but had a minor effect on SRV-X11L2, the only reporter containing a class 4 motif.

A recent analysis, which included the set of NES motifs tested here, has shown that the affinity of NESs for CRM1 linearly correlates with their nuclear export activity, for those NESs with a dissociation constant (K_d_) ranging between tens of nanomolar and tens of micromolar (Fu et al., 2018). To further investigate the relationship between CRM1 binding affinity and export activity, we plotted the previously determined K_d_ values (Fu et al., 2018) against the SRV export score of the different reporters expressed alone (Figure 3D). In line with the previous report, a clear correlation (R= −0.9225; p < 0.0001) was found between both parameters. Of note, we found that the reporter containing the high affinity superPKI NES motif (K_d_= 4 nM) exhibits higher export activity than the reporters containing NES motifs with lower affinity. This observation is in marked contrast with the reduced nuclear export activity of the superPKI NES previously reported (Fu et al., 2018). This discrepancy is most likely due to the different experimental settings used to evaluate export activity in both studies.

### Using the SRV_B/A_ assay to gain further insight into CRM1-mediated NES export. (ii) Comparing the effect of E571 and K568 mutations

A particularly intriguing and clinically relevant aspect of CRM1-mediated NES export is the role of two adjacent, electrostatically-interacting residues, namely E571 and K568 (Figure 4A). E571 mutations subtly alter nuclear export of certain NESs (García-Santisteban et al., 2016), and confer oncogenic potential to CRM1 (Taylor et al., 2019). K568 mutations (K568A or K568M), on the other hand, have been shown to allow *in vitro* binding of some “inactive NES” motifs to CRM1 by disrupting a “selectivity filter” imposed by this residue that prevents docking of structurally inadequate NES-resembling peptides (Fung et al., 2017). The relevance of this “selectivity filter” for nuclear export in a cellular setting, and the possibility that it is abrogated by cancer-related mutations in E571 remained to be investigated. Thus, we used the SRV_B/A_ assay to directly compare how the E571K and K568A mutations affect nuclear export of: (i) three “inactive NES” motifs previously characterized (COMMD1, Hxk2 and DEAF1) (Fung et al., 2017) and (ii) a subset of NES motifs from the “XPO1-cancer exportome” proteins described above.

**Figure 4.**
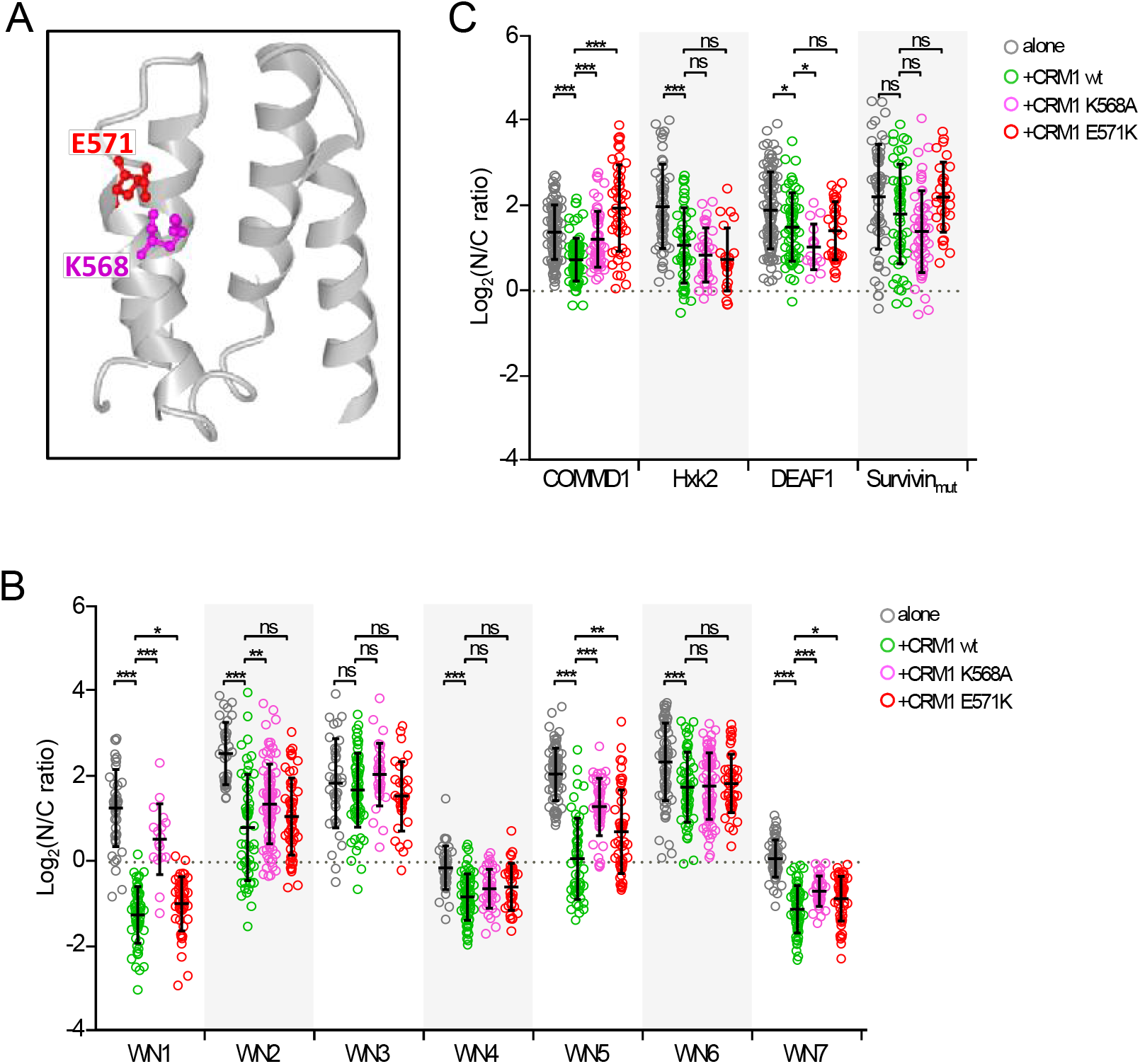
Using the SRV_B/A_ assay to compare the effect of E571 and K568 mutations. *A*. View of CRM1 NES-binding groove generated with NCBI iCn3D viewer using PDB structure 3GJX (Monecke et al., 2009). The E571 and K568 residues are highlighted using ball and stick representation, while the remaining residues are represented using ribbon style. *B*. Graph representing the nucleocytoplasmic localization of three SRV_B/A_ reporters containing “inactive” NES motifs (COMMD1, Hxk2 and DEAF1; Fung et al., 2017), plus an export defective mutant of survivin NES (Survivin_mut_), when expressed alone or when co-expressed with YFP-CRM1 (wild type, K568A or E571K). C. Graph representing the nucleocytoplasmic localization of seven SRV_B/A_ reporters containing NES motifs identified in cancer-related proteins (WN1-7) when expressed alone or when co-expressed with YFP-CRM1 (wild type, K568A or E571K). In the graphs shown in panels *B* and *C*, each circle represents a single cell where the nuclear to cytoplasmic (N/C) ratio of the fluorescent signal corresponding to the reporter was determined by image analysis using Fiji. The mean (+/− SD) is also shown. The level of statistical significance of the differences between the compared samples (Mann-Whitney U test) is indicated by the asterisks as follows: (*) p<0.05; (**) p<0.01; (***) p>0.001; ns, non-significant.

On one hand, SRV reporters containing COMMD1, Hxk2 and DEAF1 motifs were either expressed alone or co-expressed with YFP-CRM1 (wild type, K568A or E571K) in HEK293T cells. A fourth reporter, containing an export-deficient version of survivin NES mutated in two critical hydrophobic residues (García-Santisteban et al., 2016) was also included for comparison in these assays. After anti-Flag immunofluorescence, we used image analysis to quantify the intensity of the fluorescent signal of the reporters in the nucleus and the cytoplasm, and then calculated the nuclear to cytoplasmic (N/C) ratio (assessment method 3, detailed in Figure 1). As shown in Figure 4B, the localization of the four reporters was mainly nuclear (log_2_(N/C ratio)> 0) under all conditions. A minor, but statistically significant reduction in NC ratio was noted for SRV-COMMD1, SRV-Hxk2 and SRV-DEAF1 upon co-expression with wild-type CRM1, suggesting that these motifs may be NESs with extremely low activity. In comparison to the wild-type receptor, co-expression with the K568A mutant reduced export of the SRV-COMMD1 reporter, but slightly increased nuclear export of SRV-Hxk2 and SRV-DEAF1, although only the results with the later reporter reached statistical significance. These results are generally consistent with those of previous *in vitro* analyses that found increased binding of Hxk2 and DEAF1, but not COMMD1 motifs, to K568A mutant CRM1 (Fung et al., 2017). In the case of the E571K mutant, the only statistically significant effect was a reduction in the nuclear export of the SRV-COMMD1 reporter.

On the other hand, we carried out a similar analysis with a set of SRV reporters containing seven different NES motifs (WN1-WN7) identified in cancer-related proteins (Figure 4C). Consistent with the data presented in Figure 2D, the localization of all the reporters, except SRV-WN3, was significantly more cytoplasmic when co-expressed with wild-type YFP-CRM1 than when expressed alone. In comparison to the wild-type receptor, co-expression with the K568A or E571K mutants significantly reduced the export of three reporters (SRV-WN1, SRV-WN5 and SRV-WN7). As detailed in Supplementary Table 1, WN1, WN5 and WN7 motifs are three previously validated NESs in MAP kinase kinase 2 (MP2K2) (Fukuda et al., 1996), cancer susceptibility candidate gene 3 protein (CASC3) (Macchi et al., 2003) and period circadian protein homolog 1 (PER1) (Vielhaber et al., 2001), respectively. Further studies should investigate whether these proteins are aberrantly exported in E571K-mutant cells. Importantly, E571K mutation consistently led to a considerably less pronounced reduction in the nuclear export of these three reporters than K568A. Furthermore, K568A decreased nuclear export of a fourth reporter (SRV-WN2), which was efficiently exported by E571K. In summary, these results clearly indicate that the E571K mutation has a subtler effect on the nuclear export activity of CRM1 than K568A.

### Using the SRV_B/A_ assay to search for NES-harbouring micropeptides

With the progressive improvements in proteogenomics analyses, it has become apparent that the size and complexity of the cellular proteome may have been previously underestimated. Thus, there is growing evidence that a subset of RNA molecules initially annotated as non-coding may, in fact, contain short open reading frames that are translated into micropeptides, small proteins shorter than 100 amino acids in length (Yeasmin et al., 2018). Thousands of different micropeptides may be expressed in a cell, but there is still very little information on their biological function (Hartford and Lal, 2020). An important aspect of micropeptide biology that remains to be investigated is their nucleocytoplasmic localization. Given their small size, it is possible that many micropeptides can enter and exit the nucleus by passively diffusing through the nuclear pore. However, it is also possible that some micropeptides undergo active transport between the nucleus and the cytoplasm, and possess NLSs and/or NESs to interact with the nucleocytoplasmic transport machinery. To begin addressing this possibility, we decided to use the SRV_B/A_ assay to carry out a search for functional NESs in human micropeptides.

Human micropeptides were retrieved from the SmProt database, a manually curated repository of small proteins detected or predicted in eight different species (Hao et al., 2018). *In silico* prediction of putative NES motifs in the amino acid sequences of micropeptides was carried out with Wregex and NESmapper, as described above (no attempt to predict “minus” NES motifs was made in this case). Ten of the highest-ranking candidates (Supplementary Table 2) were selected and seven were successfully cloned into the SRV_B/A_ reporter for experimental testing (two representative examples are shown in Figure 5A). When expressed alone into HEK293T cells, the localization of all the reporters (determined using assessment method 1) was exclusively nuclear (N), except for SRV-MICROP-2 that showed also a faint cytoplasmic signal (N>C localization) (Figure 5B). When co-expressed with YFP-CRM1, SRV-MICROP-5 and SRV-MICROP-7 reporters fully relocated to the cytoplasm (C localization), SRV-MICROP-1 and SRV-MICROP-10 showed a minor relocation to the cytoplasm (N>C localization) and SRV-MICROP-6 and SRV-MICROP-9 remained in the nucleus. Unexpectedly, the partial cytoplasmic localization of SRV-MICROP-2 was not increased by co-expression with YFP-CRM1, suggesting that this motif may mediate CRM1-independent export or retention in the cytoplasm, rather than CRM1-mediated nuclear export. These findings identify at least two clearly active NES motifs (MICROP-5 and MICROP-7) in human micropeptides. To further confirm that these motifs are exported via CRM1, HEK293T cells co-expressing SRV-MICROP-5 or SRV-MICROP-7 with YFP-CRM1 were treated with the CRM1 inhibitor Leptomycin B (LMB). Blockade of CRM1-mediated export readily prevented cytoplasmic relocation of SRV-MICROP-7 (Figure 5C) and SRV-MICROP-5 (not shown) reporters. To our knowledge, these sequences (Figure 5D) represent the first two functional CRM1-dependent NESs identified in human micropeptides.

**Figure 5.**
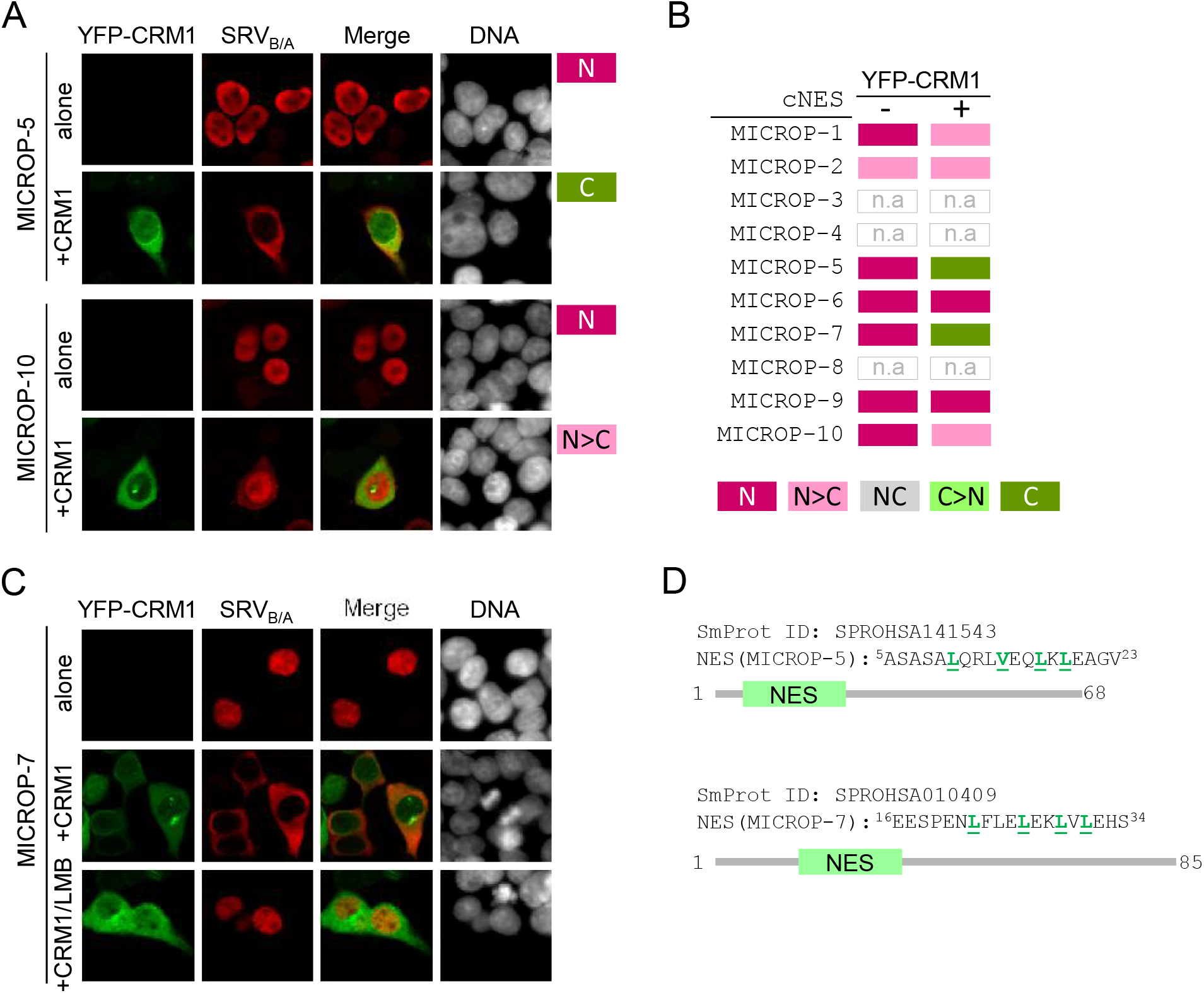
Using the SRV_B/A_ assay to search for NES-harbouring micropeptides. A. Fluorescence microscopy images showing representative examples of the localization of SRV_B/A_ reporters containing two different cNES motifs (MICROP-5 and MICROP-10, see Supplementary Table 2), when transfected alone or co-transfected with YFP-CRM1 wild type (+CRM1) into HEK293T cells. The DNA-staining dye DAPI was used to visualize the nuclei. As indicated to the right of the images, using assessment method 1 the localization of SRV-MICROP-5 was classified as N (alone) and C (+CRM1), while the localization of SRV-MICROP-10 reporter was classified as N (alone) and N>C (+CRM1). *B.* Summary of the localization of SRV_B/A_ reporters containing candidate NES motifs. The cNES ID (Supplementary Table 2) for each motif are indicated to the left. The localization of each reporter expressed alone (-) or co-expressed with YFP-CRM1 wild type (+) was assessed using method 1. Except SRV-MICROP-2, which showed a faint cytoplasmic signal (N>C localization), all reporters showed exclusively nuclear (N) localization when expressed alone. When co-expressed with YFP-CRM1, the localization of SRV-MICROP-2, SRV-MICROP-6 and SRV-MICROP-9 reporters remained invariable, SRV-MICROP-1 and SRV-MICROP-10 partially relocated to the cytoplasm and SRV-MICROP-5 and SRV-MICROP-7 fully relocated to the cytoplasm. n.a.: not assayed. *C*. Fluorescence microscopy images showing that LMB treatment blocks the cytoplasmic relocation of SRV-MICROP-7 induced by co-expression with YFP-CRM1. *D.* Schematic representation of human micropeptides SPROHSA141543 and SPROHSA010409 (SmProt IDs), showing the position of the novel NESs identified. The amino acid sequence of the NES motifs is indicated, with the hydrophobic residues that conform to the NES consensus highlighted in green.

## DISCUSSION

Our understanding of CRM1-mediated NES export derives largely from evidence obtained using structural and *in vitro* biochemical analyses with purified proteins (Dong et al., 2009a,b; Monecke et al., 2009; Güttler et al., 2010; Fox et al., 2011; Dian et al., 2013; Monecke et al., 2013; Saito and Matsuura, 2013; Fung et al., 2017). These two approaches are often combined, for example when using biochemical assays to validate predictions made from structural data (Dong et al., 2009a,b; Fox et al., 2011; Fung et al., 2017). On the other hand, cellular assays, based on the localization of reporter proteins, have been widely used to identify novel NESs or transport factors, and to test the effect of potential CRM1 inhibitors (Henderson and Eleftheriou, 2000; Fetz et al., 2009; Kehlenbach and Port, 2017), but they are not routinely used to test assumptions made from structural and *in vitro* data. In fact, a possible correlation between the affinity of NESs for CRM1 and their export activity has only recently been evaluated (Fu et al., 2018). In order to obtain a more integrative view it is important to determine how the structural and biochemical information on CRM1/NES interaction translates into NES export in the more physiological, albeit complex, cellular context.

We have recently described the SRV_B/A_, reporter (Taylor et al., 2019), a modification of the original SRV100 reporter (García-Santisteban et al., 2016), that allows evaluating how a variety of NES motifs are exported by different CRM1 variants. Here, we show how cellular assays with the SRV_B/A_ reporter can be used not only to identify novel NES motifs, but also to provide relevant information on several aspects of CRM1-mediated NES export.

All in all, we have tested 60 different (validated or candidate) NES motifs and 9 different CRM1 variants in this study. Although not every NES motif was tested for export by every CRM1 variant, our study represents, to our knowledge, the largest set of NES/CRM1 variant combinations evaluated so far. In such an extensive study, the assessment of reporter localization in a large number of samples represents a major bottleneck, unless an automated analysis platform is used. However, these sophisticated and expensive systems are not readily available to many research groups, including ours. To facilitate manual sample analysis, we decided to use three assessment methods (Figure 1C) that are increasingly time-consuming, but provide a correspondingly more detailed description of reporter localization. Thus, we reasoned that a rapid qualitative assessment of general reporter localization could be sufficient to identify functional NES motifs, but more laborious semi-quantitative methods would be required to compare the activity of the different CRM1 variants.

By combining *in silico* prediction with nuclear export assays, we report the identification of 19 novel sequence motifs with nuclear export activity in 14 proteins (SPN90, TFE3, SHIP2, PER1, SEPT6, SIR2, UBR5, FR1OP, AP2B1, IF2B, mTOR, CRTC1, CDC27 and ZO2, see Supplementary Figure 1) that belong to a putative “XPO1-cancer exportome”. Of note, this is the first attempt to specifically search not only for “classical” (“plus”) NES motifs, but also for “reverse” (“minus”) motifs. The percentage of *in silico* predicted motifs that could be experimentally confirmed was similar for both types of NES, but “plus” motifs displayed, on average, higher export activity than “minus” motifs (Supplementary Figure 1C). Thus, although the number of “minus” NESs tested so far is limited, our data suggest that these motifs are, in general, weaker than “plus” motifs.

The role of each of the identified motifs as bona-fide NESs needs to be validated, as some of these sequences might not be accessible for CRM1 binding in the context of their cognate full-length proteins. Nevertheless, our findings further support previous results suggesting that these proteins are CRM1 cargos (Kirli et al., 2015), and provide leads for further analyses of the nucleocytoplasmic localization of these important cancer-related proteins. A particularly noteworthy case is the SHIP2 protein, where two overlapping putative NES motifs were predicted, one as “plus” (WN6: 256-TGEQE<u>L</u>ESL<u>V</u>LK<u>L</u>S<u>V</u>LKDF-274) and the other as “minus” (REV5: 261-LESLV<u>L</u>K<u>L</u>SV<u>L</u>KDF<u>L</u>SGIQ-279). Both motifs displayed similar export activity in our assays. It would be interesting to apply structural analysis to establish in which orientation the SHIP2 peptide 256-279 may dock into CRM1 groove.

Testing candidate NESs in cancer-related proteins allowed us to compare the SRV_B/A_ reporter with the widely used Rev(1.4)-GFP reporter (Henderson and Eleftheriou, 2000). We found that, when using a rapid qualitative method to assess the localization of SRV_B/A_ reporters, the results obtained with both assays are well correlated. These results validate the use of the SRV_B/A_ reporter as a tool to identify novel NESs. However, it must be taken into account that, while the SRV_B/A_ assay allowed to better pinpoint differences in activity between some of the strongest NES motifs, it may miss some of the weakest NES motifs detected by the Rev(1.4)-GFP assay.

Unlike other commonly used cellular reporters, SRV_B/A_, has been applied to evaluate how different variants of CRM1 mediate export of a given NES (Taylor et al., 2019). Here, we have used SRV_B/A_-based cellular assays to test different NESs and different CRM1 variants in multiple NES/variant combinations, in an attempt to extend and complement previous observations from structural and biochemical studies.

We investigated, on one hand, how mutations in individual CRM1 groove residues affect export of a panel of well-characterized NESs motifs that belong to different classes. These NESs have variable patterns of hydrophobic residues, and structural studies have shown that they dock into the groove with different backbone conformations, varying from all helix to an almost fully extended conformation (Dong et al., 2009a; Monecke et al., 2009; Güttler et al., 2010; Fung et al., 2015; Fung et al., 2017). We did not observe a consistent relationship between NES class and how export of these motifs is affected by the different groove mutations. However, we noted that, irrespective of NES class, mutation of CRM1 residues located in the narrower part of the groove (A541, K568 and F572) consistently had a more detrimental effect on export than mutation of residues located in the wider part (I521, L525 and F561). Remarkably, mutations in K568 and F572, two residues that engage in chemically different types of interactions with the NES (main chain hydrogen bonding in the case of K568, and side chain hydrophobic interactions in the case of F572) (Dong et al., 2009b; Fung et al., 2017) reduced NES export to a similar extent. These observations suggest that interaction of the NES motif with the narrower part of CRM1 groove may be particularly relevant for efficient NES export.

Our data also allowed us to confirm the recent report that affinity for CRM1 binding (measured *in vitro*) and export activity (evaluated using a cellular assay) are linearly correlated for NES motifs across a wide range of K_d_ values (Fu et al., 2018). In fact, our observations extend these previous findings, and suggest that such a correlation is maintained even for motifs with a K_d_ below 10 nM. There was, nevertheless, a remarkable discrepancy between some of our findings and those in the previous report. We found that the SRV-superPKI reporter, containing an artificial NES motif with extremely high affinity for CRM1, was located in the cytoplasm when expressed alone in HEK293T cells. In contrast, the superPKI NES motif failed to promote efficient nuclear export of the reporter used by Fu et al. (a chimeric protein with two tandem copies of YFP and one single copy of the SV40 NLS) in HeLa cells. The different configuration of the reporters, and potential differences in the endogenous nucleocytoplasmic transport machinery of the cell lines used probably contribute to these conflicting observations, which highlights the importance of considering the influence of the experimental setting when using cellular assays to evaluate NES export activity.

On the other hand, we investigated the effect of mutations in two particularly relevant CRM1 residues, one of them (E571) recurrently mutated in human cancer, and the other (K568) reported to play a crucial role preventing docking of “inactive NES” motifs into the CRM1 groove. These residues are located in close proximity, and establish an electrostatic interaction with each other that could be abrogated by cancer-related mutations in E571. Given the close relationship between E571 and K568, we decided to directly compare the effect of mutations affecting these two residues.

We first tested CRM1 mutants K568A and E571K against three SRV_B/A_ reporters containing NES motifs previously characterized, and classified as inactive. In line with previous *in vitro* evidence showing that K568 functions as a “selectivity filter” for non-NES peptides (Fung et al., 2017), we found that two reporters containing “inactive NESs” (SRV-Hxk2 and SRV-DEAF1) were slightly better exported by a CRM1 variant carrying a K568A mutation than by the wild type receptor. The cancer-related E571K mutation, on the other hand, does not appear to abrogate this filtering effect, as it did not lead to augmented export of any of the reporters. Importantly, the minor increment in nuclear export of the SRV-Hxk2 and SRV-DEAF1 reporters afforded by the K568A mutation in our cellular assays does not reflect the markedly increased *in vitro* binding of this mutant to the Hxk2 and DEAF1 peptides reported previously (Fung et al., 2017). In this regard, it must be noted that besides abrogating the “selectivity filter”, the K568A mutation reduces binding of CRM1 to “true” NES peptides (Fung et al., 2017), and negatively impacts nuclear export activity (see our results above). These somewhat opposing effects represent a confounding factor that needs to be considered when interpreting the results of experiments with this particular CRM1 variant. Thus, while our findings are consistent with the previously described role of K568 as a “selectivity filter”, we believe that further studies are needed to better establish to what extent this residue contributes to select for “true” NES motifs.

We also compared the effect of E571K and K568A mutations against a set of SRV_B/A_ reporters containing functional NES motifs identified in cancer-related proteins. While both mutations significantly reduced the export of several of these reporters, our results clearly show that the detrimental effect of the E571K mutation is consistently less pronounced than the effect of K568A. This subtler effect on nuclear export, and our finding that E571K does not abrogate the “selectivity filter” imposed by K568, indicates that the biological consequences of E571 and K568 mutations are different. This is consistent with the fact that, while E571 mutations are highly prevalent in certain types of haematological malignancy (reviewed in Sendino et al., 2018), naturally-occurring K568 mutations have never been detected in human samples. We speculate that the subtle nature of the effects of E571K may be crucial for its oncogenic role, as mutations that more grossly disrupt CRM1-mediated export would probably be incompatible with cell survival.

Finally, after having validated the use of the SRV_B/A_ assay for NES identification on the “XPO1-cancer exportome” protein group, we used it to search for functional NES motifs within the recently uncovered “micropeptidome”, a potentially vast and still largely unexplored group of small proteins shorter than 100 amino acids in length (Yeasmin et al., 2018; Hartford and Lal, 2020). Although very few micropeptides have been characterized thus far, some of them have been shown to play a role in nuclear processes, such as DNA repair (Slavoff et al., 2014) and splicing (Huang et al., 2017), suggesting that the nucleocytoplasmic distribution of some small proteins may need to be actively regulated. Thus, our aim was to test the hypothesis that some micropeptides could represent novel CRM1 cargos. Indeed, we report here the identification of two human micropeptides (SmProt database ID SPROHSA141543 and SPROHSA010409) bearing CRM1-dependent NESs. Unlike longer proteins, where these motifs could be buried into their hydrophobic cores (Xu et al., 2012), short micropeptides (in this case, 68 and 85 amino acid long, respectively) provide a context where the identified NESs are more likely to be accessible for CRM1 binding, and thus physiologically relevant. The potential role, if any, of SPROHSA141543 and SPROHSA010409 is a mystery, as it is for the vast majority of micropeptides. It must be noted that the mere presence of these small proteins does not necessarily imply that all of them have a specific functional role. Nevertheless, our results provide the first proof-of-concept evidence, to our knowledge, that the micropeptidome can be a yet-to-be explored source of novel CRM1 cargos. Intriguingly, CRM1-binding NES motifs could also be present in some of the smallest micropeptides (15-25 amino acids in length), which raises the possibility that these micropeptides might act as “decoy NESs”, contributing to regulate CRM1-mediated nuclear export. Experiments to test this possibility are currently underway.

## ACKNOWLEDGEMENTS

We are grateful to Dr. Beric Henderson for the generous gift of the Rev(1.4)-GFP plasmid. We appreciate the technical support by the staff from the High Resolution Microscopy Facility (SGIKER-UPV/EHU).

This work was supported by grants from the Spanish Government MINECO-FEDER (SAF2014-57743-R), the Basque Country Government (IT1257-19) and the University of the Basque Country (UFI11/20), as well as a fellowship from the Basque Country Government (to MS).

The authors declare that they have no competing interests.

