## Supplementary material for "Using a simple cellular assay to map NES motifs in cancer-related proteins, gain insight into CRM1-mediated NES export, and search for NES-harbouring micropeptides"

#### SUPPLEMENTARY FIGURE LEGENDS

**Supplementary Figure 1. Analysis of candidate NES (cNES) motifs in "XPO1-cancer exportome" proteins using the Rev(1.4)-GFP nuclear export assay.**

*A.* Fluorescence microscopy images showing representative examples of the results of Rev(1.4)-GFP nuclear export assays in HeLa cells. The localization of the empty Rev(1.4)-GFP reporter (negative control) and reporters containing two different cNES motifs (WN2 and WN5) is shown. Cells were either treated (+ActD) or not (-ActD) with Actinomycin D. The DNA-staining dye DAPI was used to visualize the nuclei. The localization of the reporter in the nucleus (N), nucleus and cytoplasm (NC) or cytoplasm (C) was determined in at least 200 cells per sample. According to the percentage of cells showing each localization, the different cNESs were assigned a nuclear export score (1.4 score), as indicated in Supplementary Table 1. *B.* Schematic representation of 14 cancer-related proteins that are potential CRM1 cargos, showing the position of the 19 novel NES motifs. *C*. Graph comparing the nuclear export activity (1.4 score) of the active "plus" (n=19) and "minus" (n=6) NES motifs identified in this study. Each circle represents a single NES. The mean +/-SD is shown. The mean 1.4-score was 3.84 for "plus" motifs and 1.5 for "minus" motifs. The p value (Mann-Whitney U test) is indicated.

**Supplementary Figure 2. Raw data used to generate the heat map shown in Figure 3C.**

Graphs show the results of 126 $SRV_{B/A}$ assays testing the export activity of the indicated YFP-CRM1 variants against a panel of 14 $SRV_{B/A}$ reporters, containing previously-characterized NES motifs. For each SRV-NES/CRM1 variant combination, the localization of the reporter to the nucleus (N), nucleus and cytoplasm (NC) or cytoplasm (C) was determined in at least 200 cells. Bar colours represent the percentage of cells showing the indicated reporter localization (N, NC or C). From this percentage, a “SRV export score” was derived, as described in Methods section, and represented in the heat map shown in Figure 3C. The NES motifs tested are shown grouped according to their class.

### Supplementary Figure 1

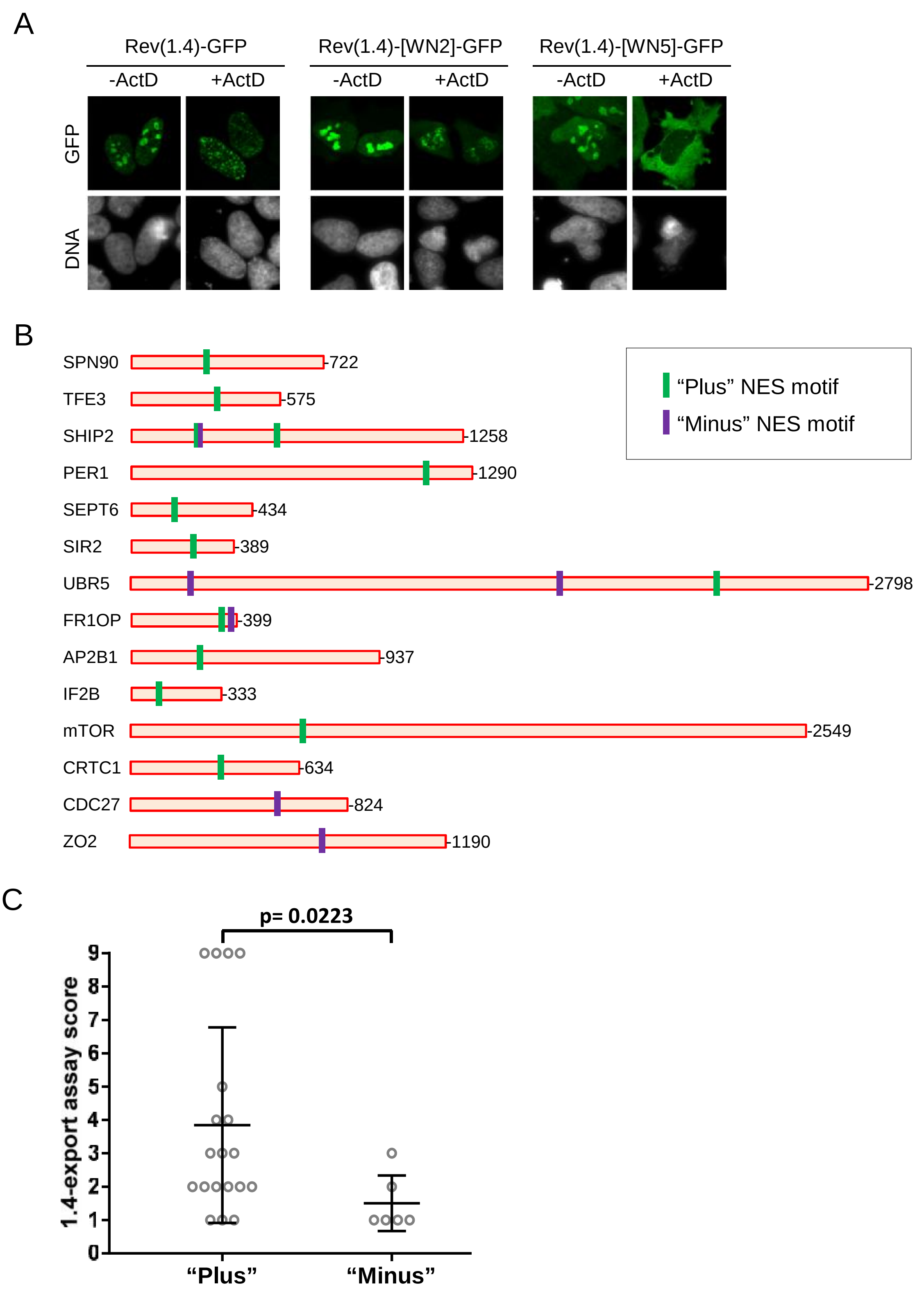

Supplementary Figure 2

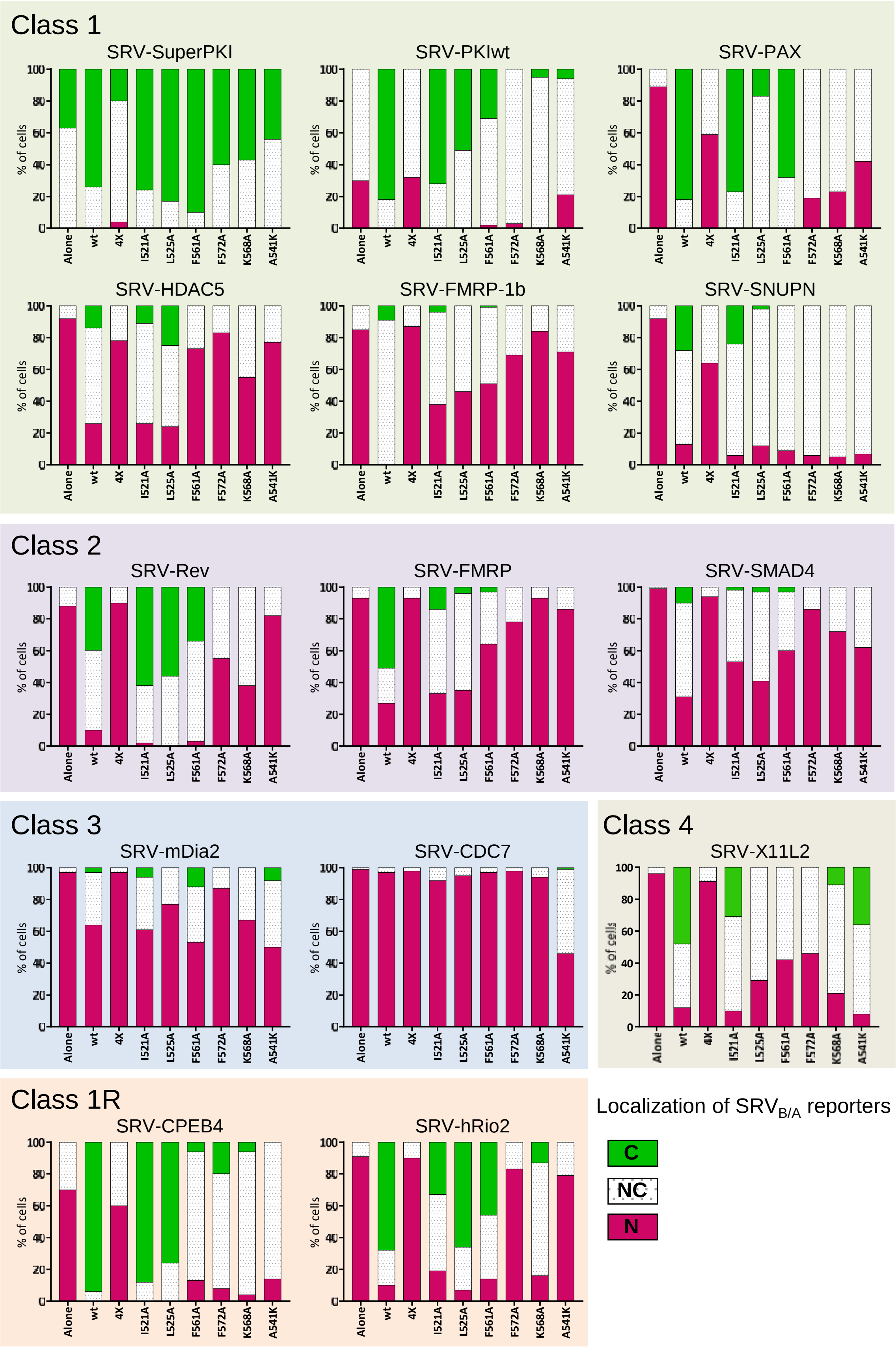

**Supplementary Table 1. Selected candidate NES motifs in "XPO1-cancer exportome" proteins predicted *in silico* and experimentally tested using the SRVB/A assay and/or the Rev(1.4)-GFP assay**

<table>
<tr><th></th><th>cNES ID</th><th>Entry [CRM1 cargo category (Kirli et al., 2015)]</th><th>cNES Position</th><th>cNES Sequence</th><th>Wregex score (quartile)</th><th>NESmapper score (quartile)</th><th>Rank</th><th>1.4 score</th><th>Previoulsy assayed? (REF)</th></tr>
<tr><td rowspan="26">Plus cNES</td><td>WN1</td><td>sp|P36507|MP2K2_HUMAN [cargo B]</td><td>33-51</td><td>---------NLVD-L-QKK-L-EE-L-E-L-DEQQK-</td><td>100 (Q1)</td><td>32,4 (Q1)</td><td>1</td><td>9</td><td>YES (Fukuda et al. 1996)</td></tr>
<tr><td>WN2</td><td>sp|Q9NZQ3|SPN90_HUMAN [cargo A]</td><td>282-300</td><td>---------SASDD-L-EA-L-GT-L-S-L-GTTEE-</td><td>100 (Q1)</td><td>13,5 (Q1)</td><td>1</td><td>3</td><td>NO</td></tr>
<tr><td>WN3</td><td>sp|P19532|TFE3_HUMAN [cargo A]</td><td>418-436</td><td>--------QANRS-L-QLR-I-QE-L-E-L-QAQI--</td><td>84,8 (Q1)</td><td>27 (Q1)</td><td>1</td><td>2</td><td>NO</td></tr>
<tr><td>WN4</td><td>sp|Q8IXJ6|SIR2_HUMAN [cargo A]</td><td>37-55</td><td>--------DMDF-L-RNL-F-SQT-L-S-L-GSQK--</td><td>74,9 (Q1)</td><td>11,2 (Q1)</td><td>1</td><td>9</td><td>YES (North and Verdin, 2007)</td></tr>
<tr><td>WN5</td><td>sp|O15234|CASC3_HUMAN [cargo B]</td><td>457-475</td><td>--------SSTSG-L-EQD-V-AQ-L-N-I-AEQN--</td><td>73,5 (Q1)</td><td>11,2 (Q1)</td><td>1</td><td>4</td><td>YES (Macchi et al., 2003)</td></tr>
<tr><td>WN6</td><td>sp|O15357|SHIP2_HUMAN [cargo B]</td><td>256-274</td><td>--------TGEQE-L-ESL-V-LK-L-S-V-LKDF--</td><td>73,5 (Q1)</td><td>8,8 (Q1)</td><td>1</td><td>3</td><td>NO</td></tr>
<tr><td>WN7</td><td>sp|O15534|PER1_HUMAN [cargo B]</td><td>483-501</td><td>-------DTDIQE-L-SEQ-I-HR-L-L-L-QPV---</td><td>70,1 (Q1)</td><td>18 (Q1)</td><td>1</td><td>9</td><td>YES (Vielhaber et al., 2001)</td></tr>
<tr><td>WN8</td><td>sp|O15357|SHIP2_HUMAN [cargo B]</td><td>625-643</td><td>--------RKEFEP-L-LR-V-DQ-L-N-L-EREK--</td><td>84,8 (Q1)</td><td>4,5 (Q2)</td><td>2</td><td>2</td><td>NO</td></tr>
<tr><td>WN9</td><td>sp|O15534|PER1_HUMAN [cargo B]</td><td>1215-1233</td><td>---------PDDP-L-FSE-L-DG-L-G-L-EPMEE-</td><td>74,9 (Q1)</td><td>6,3 (Q2)</td><td>2</td><td>1</td><td>NO</td></tr>
<tr><td>WN10</td><td>sp|Q14141-2|SEPT6_HUMAN [cargo A]</td><td>155-174</td><td>-------IAPTGHS-L-KS-L-DL-V-T-M-KKLD--</td><td>71,9 (Q1)</td><td>4,0 (Q2)</td><td>2</td><td>2</td><td>NO</td></tr>
<tr><td>WN11</td><td>sp|Q8IXJ6|SIR2_HUMAN [cargo A]</td><td>244-267</td><td>FSCMQSDFLKVDL-L-LV-M-GTS-L-Q-V-Q-----</td><td>70,1 (Q1)</td><td>4,3 (Q2)</td><td>2</td><td>2</td><td>NO</td></tr>
<tr><td>WN12</td><td>sp|O95071-2|UBR5_HUMAN [cargo A]</td><td>2206-2224</td><td>--------AEPGSI-L-TE-L-GG-F-E-V-KESK--</td><td>70,1 (Q1)</td><td>3,1 (Q2)</td><td>2</td><td>2</td><td>NO</td></tr>
<tr><td>WN13</td><td>sp|P42345|MTOR_HUMAN [cargo A]</td><td>1274-1292</td><td>-------RVSKDDW-L-EW-L-RR-L-S-L-ELL---</td><td>68,1 (Q1)</td><td>5,4 (Q2)</td><td>2</td><td>4</td><td>YES (Bachmann et al., 2006)</td></tr>
<tr><td>WN14</td><td>sp|P14635|CCNB1_HUMAN [cargo A]</td><td>138-156</td><td>---------AEED-L-CQA-F-SD-V-I-L-AVNDV-</td><td>64,9 (Q2)</td><td>18,9 (Q1)</td><td>2</td><td>5</td><td>YES (Toyoshima et al., 1998)</td></tr>
<tr><td>WN15</td><td>sp|O95684|FR1OP_HUMAN [cargo A]</td><td>352-370</td><td>---------EIS-I-GEE-I-EED-L-S-V-EIDDI-</td><td>64,9 (Q2)</td><td>15,7 (Q1)</td><td>2</td><td>1</td><td>NO</td></tr>
<tr><td>WN16</td><td>sp|P63010|AP2B1_HUMAN [cargo A]</td><td>256-274</td><td>---------VLS-A-VKV-L-MKF-L-E-L-LPKDS-</td><td>64,9 (Q2)</td><td>10,8 (Q1)</td><td>2</td><td>1</td><td>NO</td></tr>
<tr><td>WN17</td><td>sp|P20042|IF2B_HUMAN [cargo A]</td><td>89-107</td><td>--------FDIDE-A-EEG-V-KD-L-K-I-ESDV--</td><td>63,1 (Q2)</td><td>25,2 (Q1)</td><td>2</td><td>3</td><td>NO</td></tr>
<tr><td>WN18</td><td>sp|P42345|MTOR_HUMAN [cargo A]</td><td>649-668</td><td>-------VQVVADV-L-SK-L-LV-V-G-I-TDPD--</td><td>63,1 (Q2)</td><td>9,1 (Q1)</td><td>2</td><td>9</td><td>NO</td></tr>
<tr><td>WN19</td><td>sp|P11274|BCR_HUMAN [cargo A]</td><td>1091-1111</td><td>------VSGVATD-I-QA-L-KAA-F-D-V-NNKD--</td><td>63,1 (Q2)</td><td>3,7 (Q2)</td><td>2</td><td>0</td><td>NO</td></tr>
<tr><td>WN20</td><td>sp|Q14145|KEAP1_HUMAN [cargo A]</td><td>272-290</td><td>---------RCHS-L-TPN-F-LQ-M-Q-L-QKCEI-</td><td>63,1 (Q2)</td><td>3,1 (Q2)</td><td>2</td><td>0</td><td>NO</td></tr>
<tr><td>WN21</td><td>sp|Q6UUV9-3|CRTC1_HUMAN [cargo A]</td><td>329-347</td><td>----------LSP-L-SP-I-TQA-V-A-M-DALSLE</td><td>58,7 (Q2)</td><td>21,6 (Q1)</td><td>2</td><td>2</td><td>NO</td></tr>
<tr><td>WN22</td><td>sp|P15923|TFE2_HUMAN [cargo A]</td><td>566-584</td><td>--------NEAFKE-L-GR-M-CQ-L-H-L-NSEK--</td><td>58,7 (Q2)</td><td>18 (Q1)</td><td>2</td><td>0</td><td>NO</td></tr>
<tr><td>WN23</td><td>sp|Q99081|HTF4_HUMAN [cargo A]</td><td>594-612</td><td>--------NEAFKE-L-GR-M-CQ-L-H-L-KSEK--</td><td>58,7 (Q2)</td><td>18 (Q1)</td><td>2</td><td>0</td><td>NO</td></tr>
<tr><td>WN24</td><td>sp|Q12778|FOXO1_HUMAN [cargo B]</td><td>62-80</td><td>--------SAAAVS-A-DF-M-SN-L-S-L-LEES--</td><td>58,7 (Q2)</td><td>11,2 (Q1)</td><td>2</td><td>0</td><td>NO</td></tr>
<tr><td>WN25</td><td>sp|Q13492-2|PICAL_HUMAN [cargo A]</td><td>212-230</td><td>---------NEG-I-INL-L-EKY-F-D-M-KKNQC-</td><td>58,1 (Q2)</td><td>4,5 (Q2)</td><td>2</td><td>0</td><td>NO</td></tr>
<tr><td>WN26</td><td>sp|P35869|AHR_HUMAN [cargo B]</td><td>114-132</td><td>--------EGEF-L-LQA-L-NGF-V-L-V-VTTD--</td><td>58,1 (Q2)</td><td>4,5 (Q2)</td><td>2</td><td>0</td><td>NO</td></tr>
<tr><th></th><th>cNES ID</th><th>Entry [CRM1 cargo category (Kirli et al., 2015)]</th><th>cNES Position</th><th>cNES Sequence</th><th>Wregex score (quartile)</th><th>NESmapper score (quartile)</th><th>Rank</th><th>1.4 score</th><th>Previoulsy assayed? (REF)</th></tr>
<tr><td rowspan="10">Minus (reverse) cNES</td><td>REV1</td><td>sp|P25963|IKBA_HUMAN [cargo A]</td><td>267-285</td><td>-----QQLGQ-L-T-L-EN-L-QM-L-PESED-----</td><td>85,7 (Q1)</td><td>18 (Q1)</td><td>1</td><td>0</td><td>NO</td></tr>
<tr><td>REV2</td><td>sp|Q16204|CCDC6_HUMAN [cargo A]</td><td>297-315</td><td>-----MREEN-L-R-L-QRK-L-QRE-M-ERR-----</td><td>85,7 (Q1)</td><td>16,2 (Q1)</td><td>1</td><td>0</td><td>NO</td></tr>
<tr><td>REV3</td><td>sp|P19484|TFEB_HUMAN [cargo A]</td><td>431-449</td><td>-----KDLDL-M-L-L-DDS-L-LPL-A-SDP-----</td><td>76,2 (Q1)</td><td>36 (Q1)</td><td>1</td><td>0</td><td>NO</td></tr>
<tr><td>REV4</td><td>sp|O95071-2|UBR5_HUMAN [cargo A]</td><td>209-227</td><td>-----LQRTN-L-D-V-NLA-V-NN-L-LSRD-----</td><td>74,9 (Q1)</td><td>9 (Q1)</td><td>1</td><td>2</td><td>NO</td></tr>
<tr><td>REV5</td><td>sp|O15357|SHIP2_HUMAN [cargo B]</td><td>261-279</td><td>-----LESLV-L-K-L-SV-L-KDF-L-SGIQ-----</td><td>73,5 (Q1)</td><td>49,5 (Q1)</td><td>1</td><td>3</td><td>NO</td></tr>
<tr><td>REV6</td><td>sp|P30260|CDC27_HUMAN [cargo A]</td><td>545-563</td><td>-----HLQKD-V-A-L-SV-L-SKD-L-TDMD-----</td><td>73,5 (Q1)</td><td>16,2 (Q1)</td><td>1</td><td>1</td><td>NO</td></tr>
<tr><td>REV7</td><td>sp|O95071-2|UBR5_HUMAN [cargo A]</td><td>1607-1625</td><td>-----EDGSD-M-E-L-DL-L-AA-A-ETESD-----</td><td>71,9 (Q1)</td><td>13,5 (Q1)</td><td>1</td><td>1</td><td>NO</td></tr>
<tr><td>REV8</td><td>sp|Q9UDY2-3|ZO2_HUMAN [cargo A]</td><td>715-733</td><td>------PIAD-I-A-M-EK-L-ANE-L-PDWFQ----</td><td>70,1 (Q1)</td><td>17,6 (Q1)</td><td>1</td><td>1</td><td>NO</td></tr>
<tr><td>REV9</td><td>sp|Q92997-2|DVL3_HUMAN [cargo A]</td><td>24-42</td><td>------PAER-V-T-L-AD-F-KGV-L-QRPSY----</td><td>70,1 (Q1)</td><td>12,6 (Q1)</td><td>1</td><td>0</td><td>NO</td></tr>
<tr><td>REV10</td><td>sp|O95684|FR1OP_HUMAN [cargo A]</td><td>379-397</td><td>------LTQD-L-T-V-SQ-L-SDV-A-DYLED----</td><td>70,1 (Q1)</td><td>10 (Q1)</td><td>1</td><td>1</td><td>NO</td></tr>
</table>

**Supplementary Table 2. Selected in silico predicted candidate NES motifs in human micropeptides**

| cNES ID | SmProt peptide ID and size (aa) | cNES Position | Sequence | Wregex score (quartile) | NESmapper score (quartile) | Rank |
|---|---|---|---|---|---|---|
| MICROP-1 | SPROHSA011142 (96) | 13-31 | ----KKEE-L-LKQ-L-DD-L-K-V-ELSQL | 86,5 (Q1) | 22,6 (Q1) | 1 |
| | SPROHSA011145 (47) | 13-31 | | | | |
| MICROP-2 | SPROHSA018908 (84) | 57-75 | ----IRDR-L-PVN-V-RE-L-S-L-DDPEV | 86,5 (Q1) | 11,7 (Q1) | 1 |
| MICROP-3 | SPROHSA012652 (70) | 29-47 | ----GLDD-L-DVA-L-SN-L-E-V-KLEGS | 85,7 (Q1) | 34,2 (Q1) | 1 |
| MICROP-4 | SPROHSA141226 (78) | 40-58 | ---DGTSD-L-PLK-L-EA-L-S-V-KEDA- | 85,7 (Q1) | 20,2 (Q1) | 1 |
| | SPROHSA141826 (85) | 47-65 | | | | |
| MICROP-5 | SPROHSA141543 (68) | 5-23 | ---ASASA-L-QRL-V-EQ-L-K-L-EAGV- | 83,7 (Q1) | 30,6 (Q1) | 1 |
| MICROP-6 | SPROHSA011811 (57) | 30-48 | --SHYHET-L-GEA-L-QG-V-E-L-EFS-- | 83,7 (Q1) | 17,6 (Q1) | 1 |
| MICROP-7 | SPROHSA010409 (85) | 16-34 | --EESPEN-L-FLE-L-EK-L-V-L-EHS-- | 83,7 (Q1) | 13,8 (Q1) | 1 |
| MICROP-8 | SPROHSA009911 (99) | 78-96 | --RMSKEE-L-RAK-L-SE-F-K-L-ETR-- | 83,7 (Q1) | 12,6 (Q1) | 1 |
| MICROP-9 | SPROHSA020870 (100) | 46-64 | --LSKCGEE-L-GR-L-KL-V-L-L-ELN-- | 82,5 (Q1) | 16,3 (Q1) | 1 |
| MICROP-10 | SPROHSA180177 (93) | 49-67 | AKIKLLTKE-L-SV-L-KD-L-F-L-E---- | 81,1 (Q1) | 11,2 (Q1) | 1 |
| | SPROHSA180747 (93) | 49-67 | | | | |
| | SPROHSA181614 (93) | 49-67 | | | | |
